# Variant Set Distillation

**DOI:** 10.1101/2024.12.06.627210

**Authors:** Ryan Christ, Chul Joo Kang, Louis J.M. Aslett, Daniel Lam, Maria Faelth Savitski, Nathan Stitziel, David Steinsaltz, Ira Hall

## Abstract

Allelic heterogeneity – the presence of multiple causal variants at a given locus – has been widely observed across human traits. Combining the association signals across these distinct causal variants at a given locus presents an opportunity for empowering gene discovery. This opportunity is growing with the increasing population diversity and sequencing depth of emerging genomic datasets. However, the rapidly increasing number of null (non-causal) variants within these datasets makes leveraging allelic heterogeneity increasingly difficult for existing testing approaches. We recently-proposed a general theoretical framework for sparse signal problems, Stable Distillation (SD). Here we present a SD-based method `vsdistill`, which overcomes several major shortcomings in the simple SD procedures we initially proposed and introduces many innovations aimed at maximizing power in the context of genomics. We show via simulations that `vsdistill` provides a significant power boost over the popular STAAR method. `vsdistill` is available in our new R package `gdistill`, with core routines implemented in C. We also show our method scales readily to large datasets by performing an association analysis with height in the UK Biobank.

## 1 Introduction

While genome-wide association studies (GWASs) have successfully mapped common traits and diseases to many causal genes across the genome, recent heritability studies estimate that there are many more causal genomic loci that have yet to be discovered because the causal variants they harbor have weak effects or are rare in sequenced human populations [31]. Several variant set and gene testing methods have been successful at uncovering previously unidentified causal loci by leveraging allelic heterogeneity (AH): the presence of more than one independent causal variant at a given locus [32, 19, 30].

Growing evidence suggests that AH is common for human traits [17, 10]. A recent *in vitro* study of identified associations estimated that 10% to 20% of expression quantitative trait loci (eQTLs) have multiple causal regulatory variants [1]. These results underscore the potential role of AH in identifying genes contributing to disease risk [23, 8, 31]. The opportunity for methods that can leverage AH to identify overlooked associations will only increase with the sample size, population diversity, and sequencing depth of genomic datasets. However, all three of these factors introduce more variants to a given dataset, most of which will be non-causal (null). This increases the multiple testing burden at each locus, thereby diminishing statistical power.

Maintaining power in this sparse signal setting – where only few of many variants in a given gene or variant set are causal – is difficult for any statistical method due to the complex dependence structure (LD) among nearby variants. Popular early methods for dealing with this dependence, such as the Sequential Kernel Association Test (SKAT) [32], struggled in this context because they were based on quadratic form test statistics, which have provably little power against sparse signals [2]. The Cauchy Combination Test (CCT) [21] opened a second approach that has been taken by popular current methods, notably ACAT-O [22] and STAAR [20]. Rather than model the dependency between nearby variants, the CCT simply takes marginal p-values calculated independently for each variant and applies a version of Fisher’s classic p-value combination method that is robust to some forms of dependency among the p-values, yielding an approximately calibrated overall p-value.

However, the ability of the CCT to combine independent association signals across variants is limited. When faced with a set of variants where two of those variants have very small p-values, an ideal statistical method would combine those two p-values if the variants were uncorrelated (unlinked) and effectively discard one of those p-values as redundant if the variants were very correlated (strongly linked). Since the CCT does not have access to the correlation structure among the variants, it cannot distinguish between these two cases. Thus, to maintain calibration, the CCT must over-penalize variant sets where there are two independent signals, leading to a loss in power. This argument extends to the case of more than two causal variants. Indeed, the p-value returned by the CCT cannot be more significant than the smallest p-value in the initial set. Barnett et al. proposed a robustified version of a popular outlier test, Tukey’s higher criticism [28, 14], for testing variant sets [4]. However, that approach has not been used much since, similar to the CCT, it cannot distinguish between observations where two p-values are driven by correlated variants or independent variants, limiting its potential power gains.

We recently proposed a new statistical framework, Stable Distillation (SD), which effectively decouples the dependency structure among predictors [7]. When applied to variant set testing, this approach assigns an independent p-value to each variant under the null hypothesis that there is no association between the variants and the phenotype of interest. This allows us to test the null hypothesis using outlier tests that are optimized for cases where, roughly speaking, a handful of p-values are significant. Based on the SD framework, here we present a new variant set testing method built around a new procedure we call Helical Distillation.

In spirit, the stochastic process underlying Helical Distillation mimics the behavior of classic forward stepwise regression [15], sometimes referred to as sequential conditional analysis in the genomics literature, where a practitioner iterates a two step procedure: a) running single marker testing on a set of variants and then b) adding the most significant variant to the regression model. This simple model selection approach, with the addition of a variable rejection step, is at the heart of conditional and joint (COJO) analysis [34], which has become popular for running genome-wide association studies where one starts with all variants in the genome (see the implementation provided in GCTA for details [33]). Obtaining an overall p-value against the global null hypothesis that there is no association between the hypothesis vectors and the phenotype based on forward stepwise regression is difficult because the p-values returned by simple forward stepwise regression do not account for the “winner’s curse” – the sequential selection of the most associated hypotheses – and they are not independent. Crucially, the p-values returned by Helical Distillation account for the “winner’s curse” and are guaranteed to be mutually independent under the global null hypothesis. This allows us to use Helical Distillation as the basis for developing powerful tests against the global null hypothesis. Due to the close similarity between forward stepwise regression and linear model boosting and LASSO regression [15] – both fundamental machine learning tools for tackling sparse-signal problems – Helical Distillation provides a new solution to an important and ongoing challenge in the statistics literature. As we describe further in Section 4, Helical Distillation makes several improvements on the simple SD procedures we initially presented in [7]. Most notably, Helical Distillation has substantially improved power in the case of heterogeneous effect sizes across active predictors (causal variants).

Helical Distillation is just one of five stages in the genotype-to-phenotype variant set distillation procedure, `vsdistill` . It is implemented in our new R-package, `gdistill`, with core routines written in C. `gdistill` will be made publicly available at `ryanchrist.r-universe.dev` shortly. Please to request access to a pre-release version. Like existing variant set testing methods, `vsdistill` can admit prior probabilities that each variant is causal. That prior information is seamlessly propagated through all stages of the procedure, making it easy to incorporate variant impact predictions from emerging methods such as AlphaMissense [5]. `vsdistill` can also incorporate estimates of rare variant burden or gene function such as those provided by DeepRVAT [9].

In extensive simulations, we demonstrate that `vsdistill` yields substantial power gains over a leading variant-set testing approach STAAR. We illustrate the scalability of `vsdistill` by appyling it to height in whole-exome-sequencing (WES) data from the UK Biobank [3].

## 2 Results

### 2.1 A distillation-based VST framework

As depicted in Figure 1, our variant set distillation procedure, `vsdistill`, is a five-stage genotype-to-phenotype testing procedure. The first stage, hypothesis generation, begins with a matrix of genotypes *G*. In this stage, the genotypes are used to generate potential predictors of the phenotype *Y* . By default, each genotype is then converted into three “hypothesis” vectors, each corresponding to a different inheritance model: additive, dominant, or recessive. Optionally, a prior probability *θ* that a given genotype affects the phenotype under a given inheritance model may be assigned to each hypothesis vector. By default, each hypothesis is given equal prior weight. See Section 4.2 for details.

**Figure 1.**
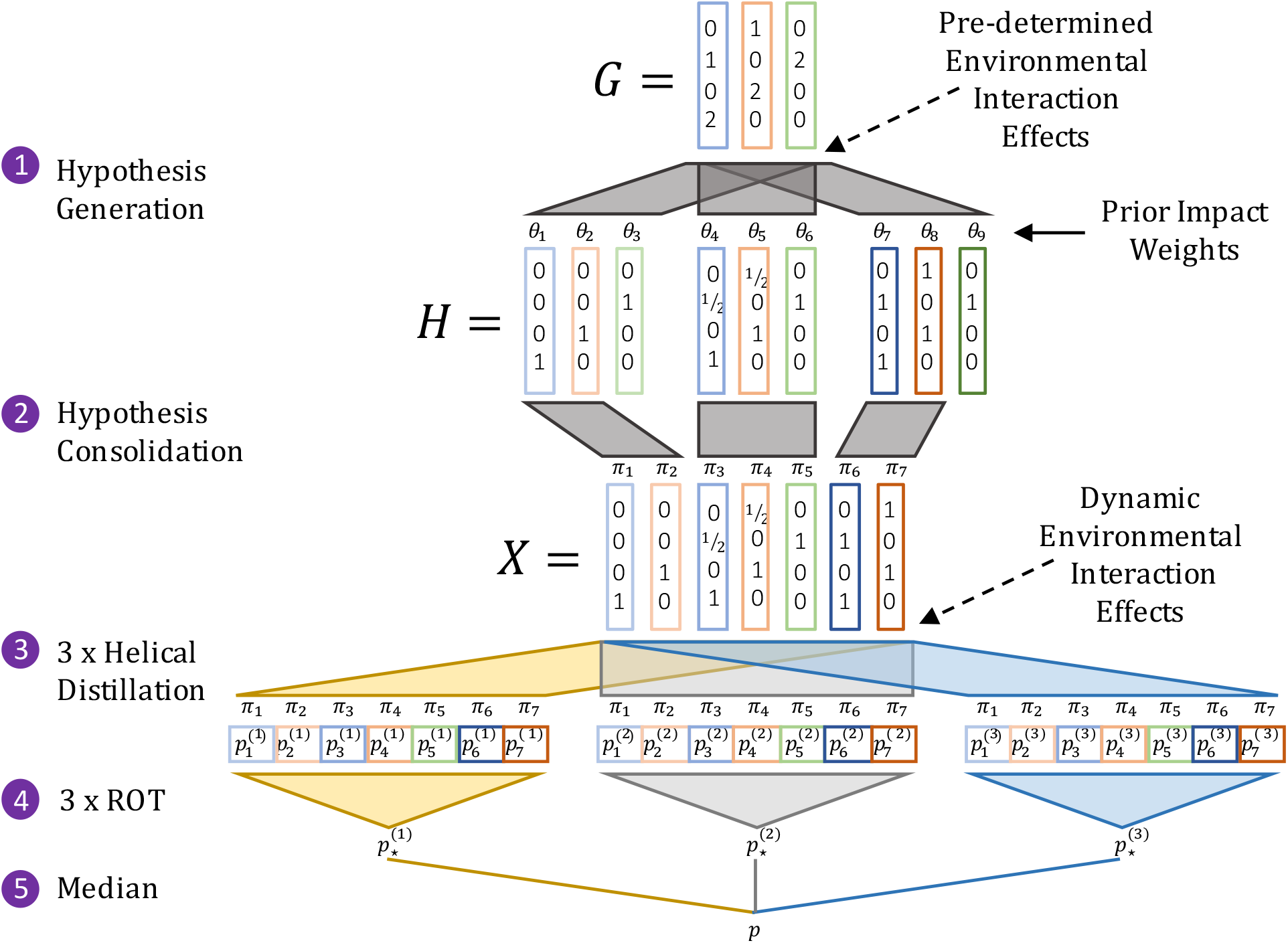
Variant Set Distillation (vsdistill): Our 5-stage procedure starts with a matrix of *g* genotypes 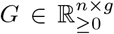. In Stage 1, Hypothesis Generation, we generate a matrix of hypothesis vectors *H∈* ℝ ^*n*×*h*^ From *G*. By default, each genotype vector is mapped to three hypothesis vectors, each reflecting a different inheritance model (recessive, additive, and dominant). Prior variant impact prediction probabilities, *θ* may be included at this stage. In Stage 2, redundant hypotheses are pruned, yielding a consolidated matrix of predictors *X* ∈ ℝ ^*n*×*p*^ with updated weights 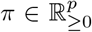 based on the initial *θ*. In Stage 3, we apply Helical Distillation to *X* and the phenotype vector *Y* in three separate replicates, yielding three vectors of p-values *p*^(1)^, *p*^(2)^, *p*^(3)^ ∈ [0, 1]^*p*^. In Stage 4, the Rényi Outlier Test (ROT) maps each vector of p-values to a single p-value using prior weights *π*. In Stage 5, the median p-value among these final three p-values is multiplied by 3/2 and returned as our final `vsdistill` p-value *p*.

The second stage, hypothesis consolidation, deploys a greedy clustering algorithm to obtain a reduced set of representative hypothesis vectors and corresponding weights. This greedy algorithm is reminiscent of LD pruning but appropriately accounts for background covariates and the prior probabilities assigned to each genotype. The weight *π* assigned to each representative hypothesis vector is taken to be proportional to the prior probability of that vector affecting the phenotype. Each *π* is calculated based on the prior probabilities *θ* of the original hypotheses that the representative vector represents. Consolidating hypotheses increases our statistical power by reducing the effective multiple testing burden. See Section 4.3 for details.

Stage three, Helical Distillation, is the core of our approach. It iteratively tests the representative hypothesis vectors for association with the phenotype vector, and assigns a p-value to each representative hypothesis while prioritizing hypotheses with a stronger association with the phenotype and higher prior probability. See Section 4.4 for details.

In stage four, we use the Rényi Outlier Test (ROT) to combine the p-value assigned to each representative vector hypothesis into a single association p-value [6]. Intuitively, this approach applies a lower multiple testing burden to association signals arising from representative hypotheses with higher prior probabilities. Since the Helical Distillation step is inherently stochastic, the p-values it returns can change between runs.

This makes the downstream single p-value returned by the ROT stochastic. While our simulations show very little variation in these p-values across runs when an association signal is present, to reduce this stochasticity, we re-run the third and fourth stages of `vsdistill` three times and take the middle p-value. More explicitly, in Stage 5, we multiply the second smallest of the three p-values by 3*/*2 to obtain our final `vsdistill` p-value. This follows from a generalization of the Bonferroni method to order statistics of dependent p-values [25].

Although not explicitly shown in Figure 1, our approach can account for background covariates and be used to boost non-additive effects captured by loss-of-function predictions, burden testing, or more general gene impairment scores [9]. See Section 4.1 for details.

### 2.2 Type-I Error Control

In order to confirm the calibration of `vsdistill` empirically, we simulated 5 thousand independent genomic datasets, each consisting of 30 thousand samples of a 1 Mb chromosome. See Section 4.5 for more details. For each dataset, we simulated 1 thousand independent phenotype vectors assuming no causal variants. This yielded a total of 5 million independent phenotype vectors. Each vector was simulated under the null model *Y* ∼ *N* (*Aα, I*_*n*_). Our definition of the background covariates *A, α*, and other details of our phenotype simulation approach are in Section 4.6. We tested each phenotype vector for association with genotypes within the central 10 kb window of corresponding chromosome using `vsdistill` . The resulting Q-Q plot (Figure 4, Supplemental Information) confirms that `vsdistill` is well calibrated under the null hypothesis.

### 2.3 Power

We compared `vsdistill` to ACAT-O (STAAR without prior weights) and standard SMT across a variety of genetic architectures. We considered *a* = 3, 9, or 15 causal variants based on the number of independent causal alleles that were observed in the large follow-up study of GWAS hits by Abell et al. (see Figure 4b of [1]). In each simulation, causal variants were randomly assigned from among those with derived allele frequency less than 0.01 within a 10 kb causal window in the center of each simulated 1 Mb segment.

Under each genetic architecture, we estimated power as a function of the underlying total association signal strength: the − log_10_ p-value that one would obtain by testing the simulated phenotype *Y* with an oracle ANOVA model that “knows” which are the active predictors and targets only those for testing. In our first batch of simulations, following [7], we used the *QR*-decomposition to ensure that the total association signal was evenly split among the causal variants in every simulation. In other words, we ensured that the *observed* contribution of each causal variant to the total association signal was essentially equal for each simulated *Y*. We refer to these as our equal “observed” signal simulations. We then replicated our power simulations in a more traditional way, simply setting the effect size *β* equal among the causal variants (after variance normalizing each genotype). This ensured that the *expected* contribution of each causal variant to the total association signal was essentially equal for each simulated *Y* in our second batch of simulations. We refer to these as our equal “expected” signal simulations.

For any particular *Y* among the equal “expected” signal simulations, the observed contribution of each causal variant to the overall association signal can vary by orders of magnitude, with one causal variant typically standing out as the main driver of the observed association signal. This means that equal “expected” signal simulations should be understood as being somewhere in between the situation where there is only one causal variant and the situation where there are multiple causal variants equally contributing to the association signal. Please see Section 4.6 for more details of our phenotype simulation approach.

We compared three variant-set tests: `vsdistill`, STAAR, and the oracle ANOVA method that “knows” the active predictors (provided as a benchmark). To calculate power for these variant-set tests, we counted our causal region as “discovered” if a test returns a p-value less than 0.05*/*20, 000. We also compared to standard SMT; we counted our causal region as “discovered” by SMT if any SMT p-value falls below 10^−8.5^ threshold *anywhere* along the entire 1 Mb region. This definition reflects how new associations are discovered in practice and provides a relatively strict benchmark.

Each point of the resulting power curves was estimated via 2 thousand independent samples: we simulated 10 independent phenotype vectors for each of 200 independent genomic datasets, each consisting of a 1 Mb chromosome sampled for 30 thousand individuals. For an example, one set of power curves, for the case of 9 causal variants with equal “expected” signals, is shown in Figure 2. The rest are shown in Supplemental Figures 5, 6, 7,8,9. We summarize all of the curves with the estimated minimum signal strength required to achieve 80% power (lower is better) in Figure 2.

**Figure 2.**
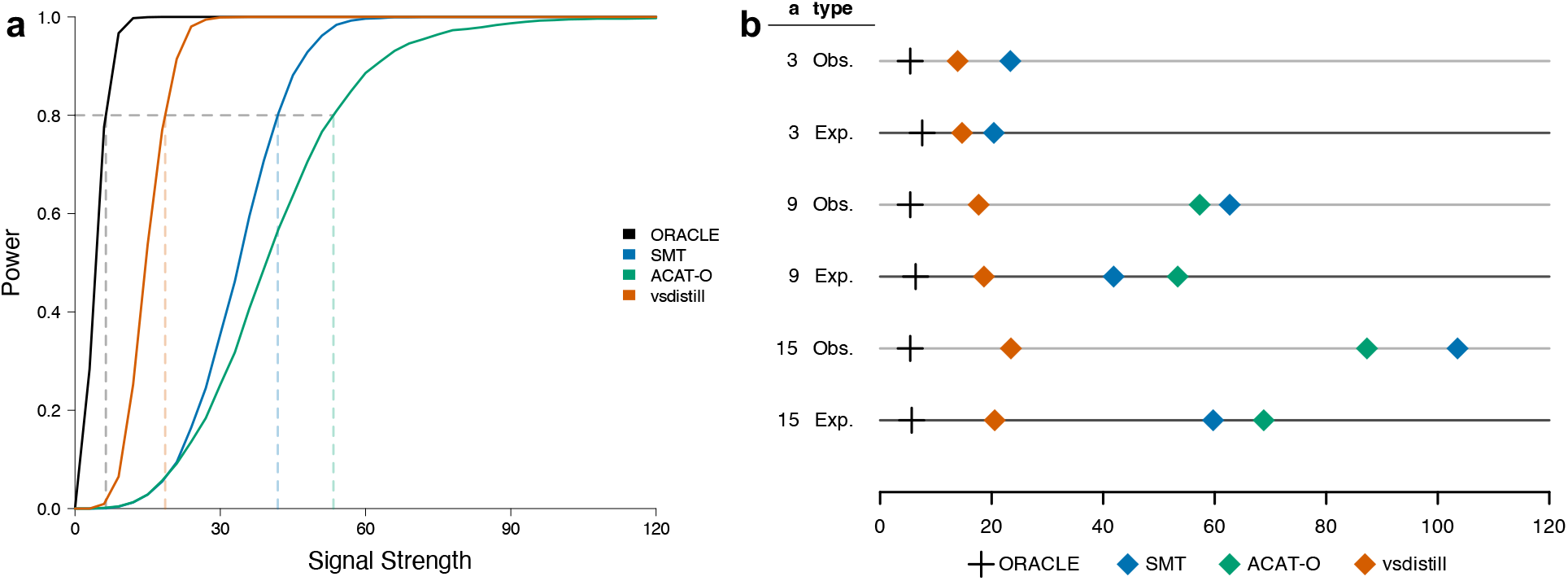
Power Simulations. **a)** Power curves for the case of 9 causal variants with equal “expected” effects as a function of total signal strength *s* (see Section 4.6). Dotted lines denote the signal strength required to achieve 80% power for each method. **b)** Dotplot showing the signal strength required to achieve 80% across methods and simulation set ups. The first column, **a**, reports the number of causal variants. The second column, **type**, reports whether the simulations targeted equal “expected” or equal “observed” effects across the causal variants. In cases where a method never achieved 80% power over then range of simulated signal strengths, a dot (diamond) for that method was not plotted.

Our power simulations demonstrate the substantial improvements in power provided by `vsdistill`, with more impressive improvements when a given association signal is divided among more causal variants. This power gain is even more notable given the fact that only additive effects were simulated here, while `vsdistill` tested all three inheritance models – dominant, recessive, and additive – for every variant, yielding a much higher multiple testing burden. In contrast, here ACAT-O and SMT only test the additive model.

### 2.4 Application to Height in the UK Biobank

In order to demonstrate the scalability and effectiveness of `vsdistill` in real genomic data, we screened 18,414 genes for association with standing height using WES data from 323,529 samples with European ancestry in the UK Biobank. As shown in Figure 1, `vsdistill` returns three replicate p-values *p*^(1)^, *p*^(2)^, *p*^(3)^ at the end of Step 4. We concatenated these three p-values across all genes and applied genomic control (*λ* = 1.67) to the full collection of p-values [13]. This yielded the Q-Q plot in Figure 3a. A staggering 21% of the genome is estimated to affect height [35], and that estimate is conservative since it is just based on common variation. Thus we expect a well-calibrated Q-Q plot for this experiment to show alignment between expected and observed quantiles from 0 up to roughly − log_10_(0.21) ≈ 0.62, which is what we observe in Figure 3a.

**Figure 3.**
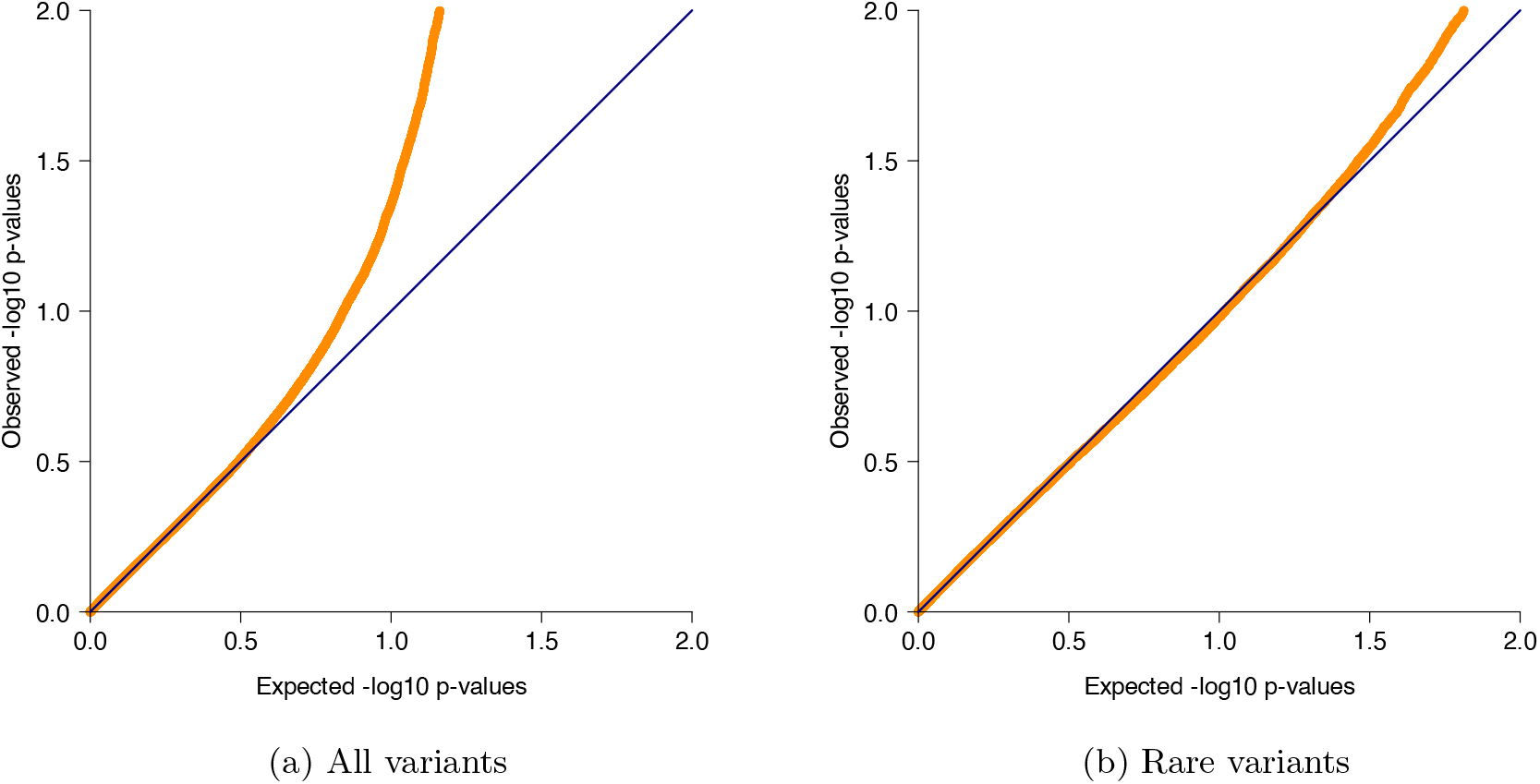
Genome-wide calibration of vsdistill on UK Biobank exonic variants: Q-Q plots of `vsdistill` − log_10_ p-values after applying genomic control. These plots include three p-values per gene because we pool together and calibrate the three replicate p-values returned by the ROT (Step 4) before taking the median (Step 5). The Q-Q plots are zoomed into the bottom left corner to show calibration in the body of the distribution.

**Figure 4.**
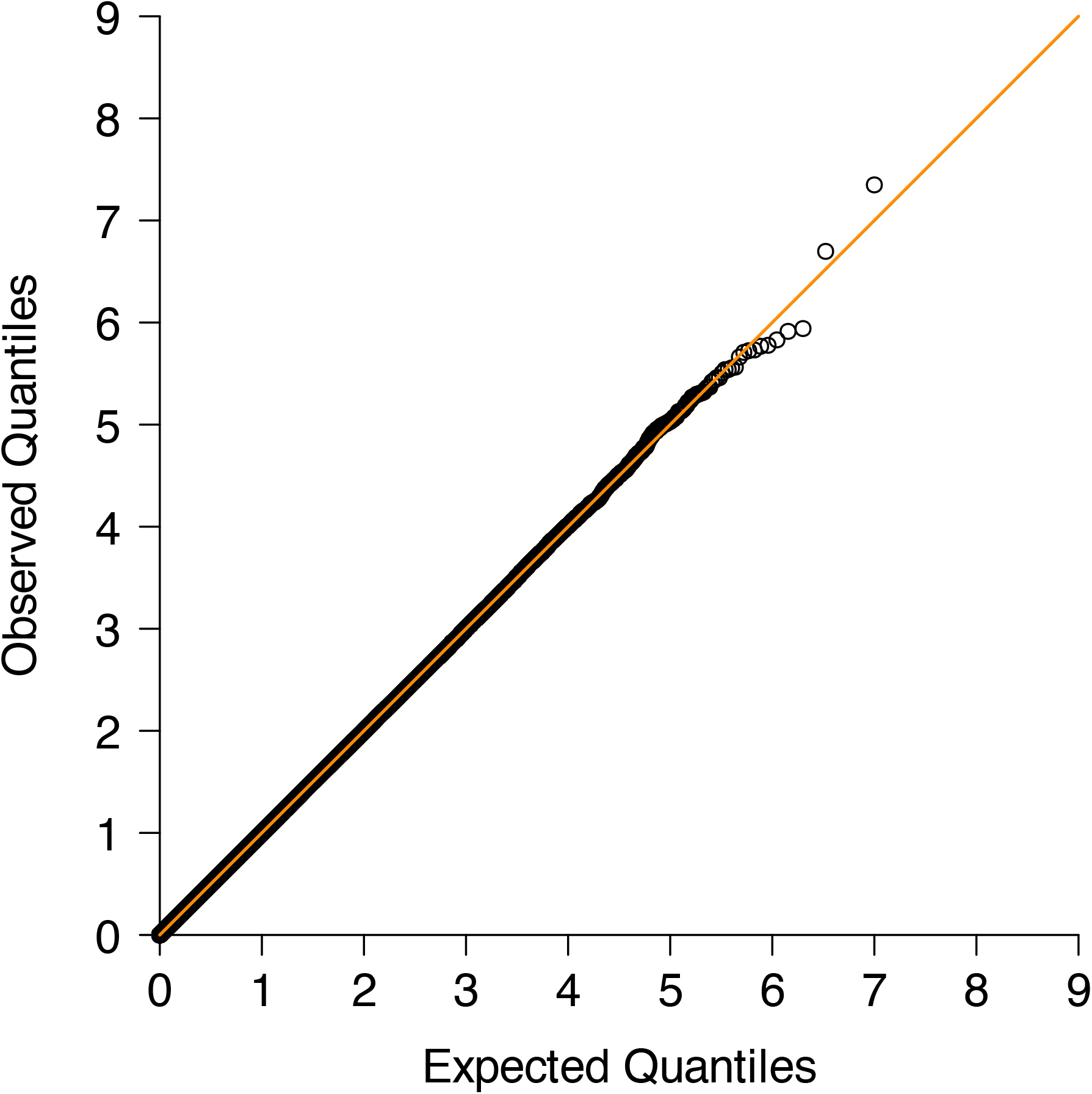
Q-Q plot of − log_10_ p-values returned by `vsdistill` across 5 million null simulations.

**Figure 5.**
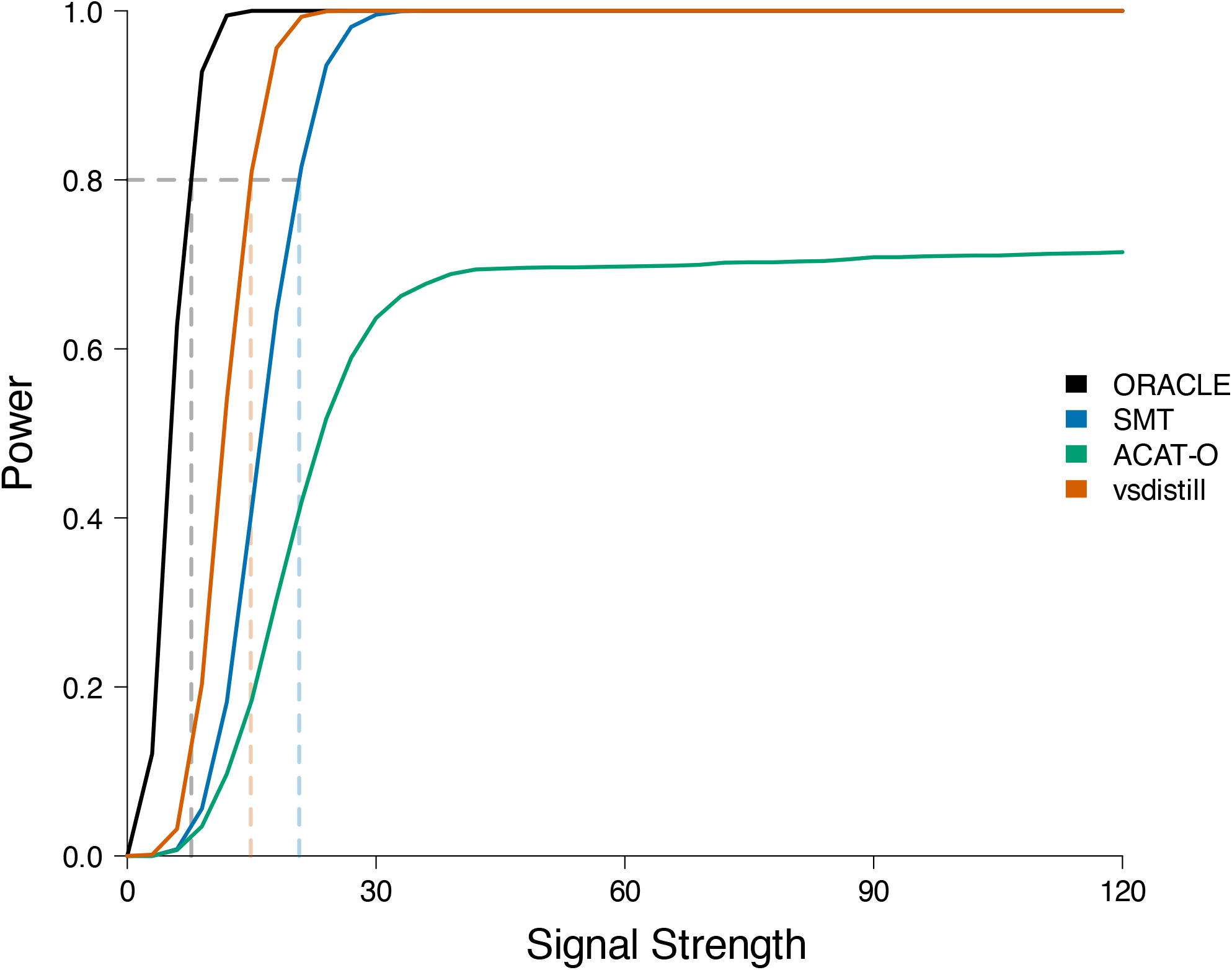
Power comparison between `vsdistill`, SMT, and ACAT-O in simulations where there are 3 causal variants with equal “expected” effects. The power an oracle ANOVA gene test that knows the causal variants is also included as a benchmark.

**Figure 6.**
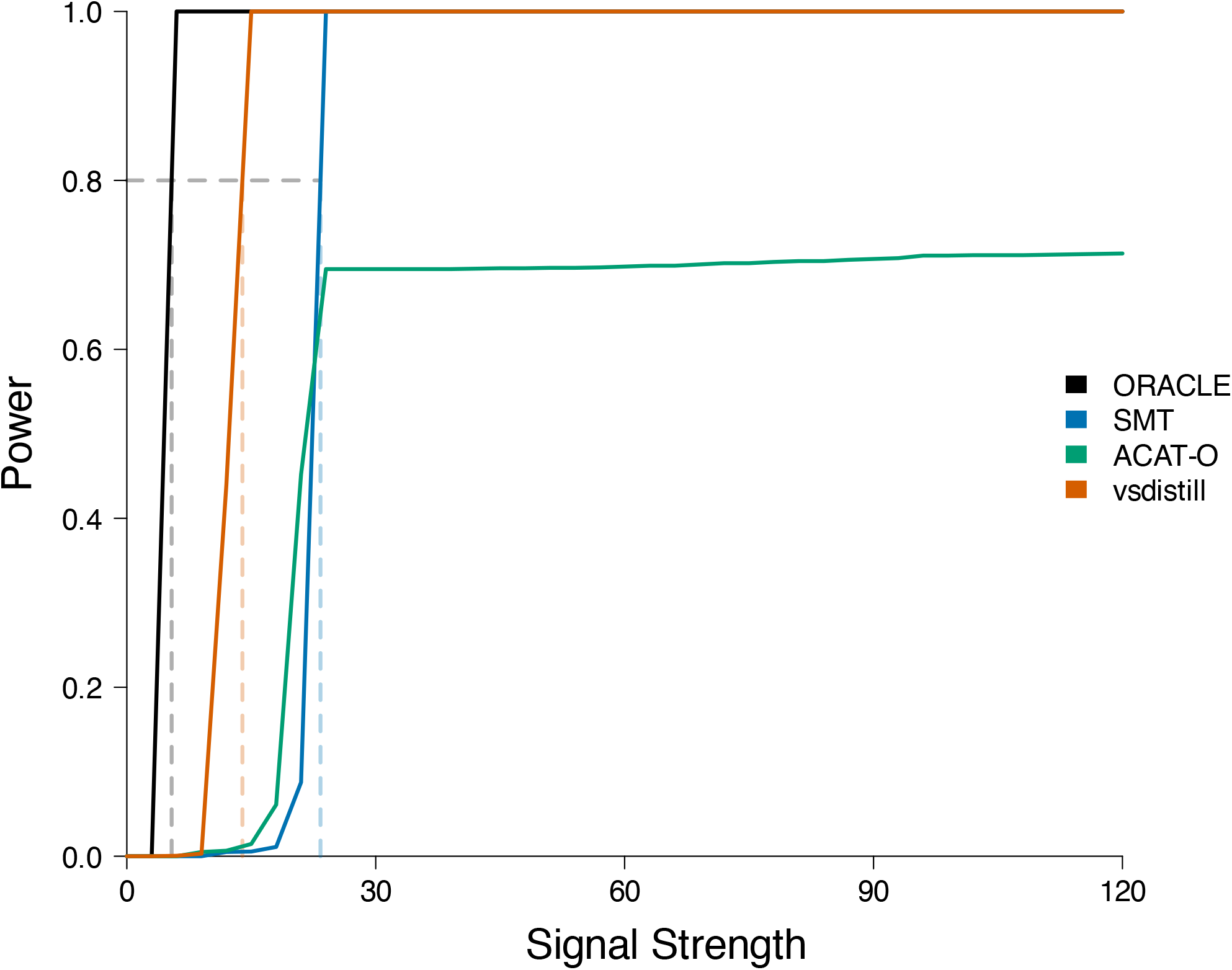
Power comparison between vsdistill, SMT, and ACAT-O in simulations where there are 3 causal variants with equal “obseved” effects. The power an oracle ANOVA gene test that knows the causal variants is also included as a benchmark.

**Figure 7.**
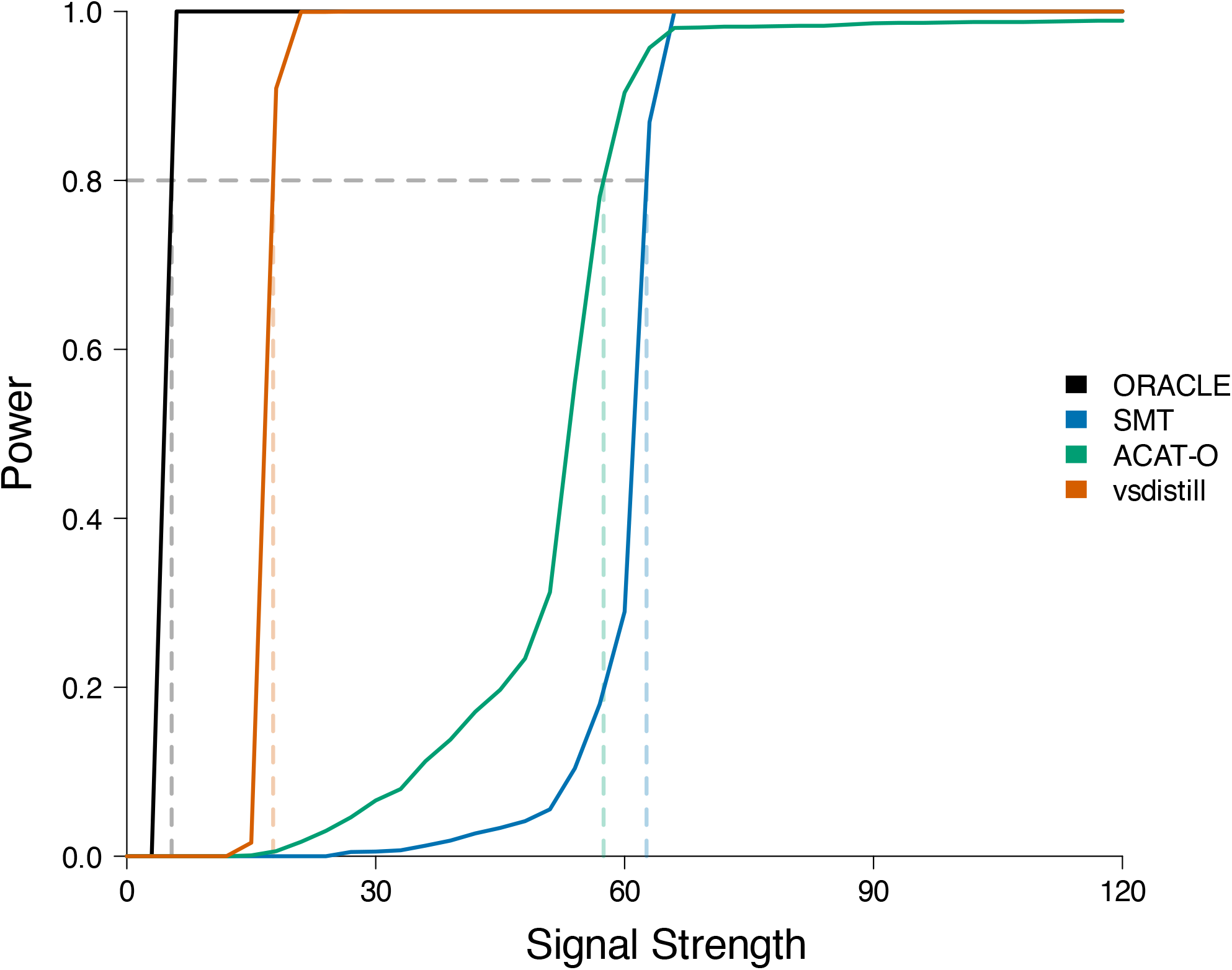
Power comparison between vsdistill, SMT, and ACAT-O in simulations where there are 9 causal variants with equal “obseved” effects. The power an oracle ANOVA gene test that knows the causal variants is also included as a benchmark.

**Figure 8.**
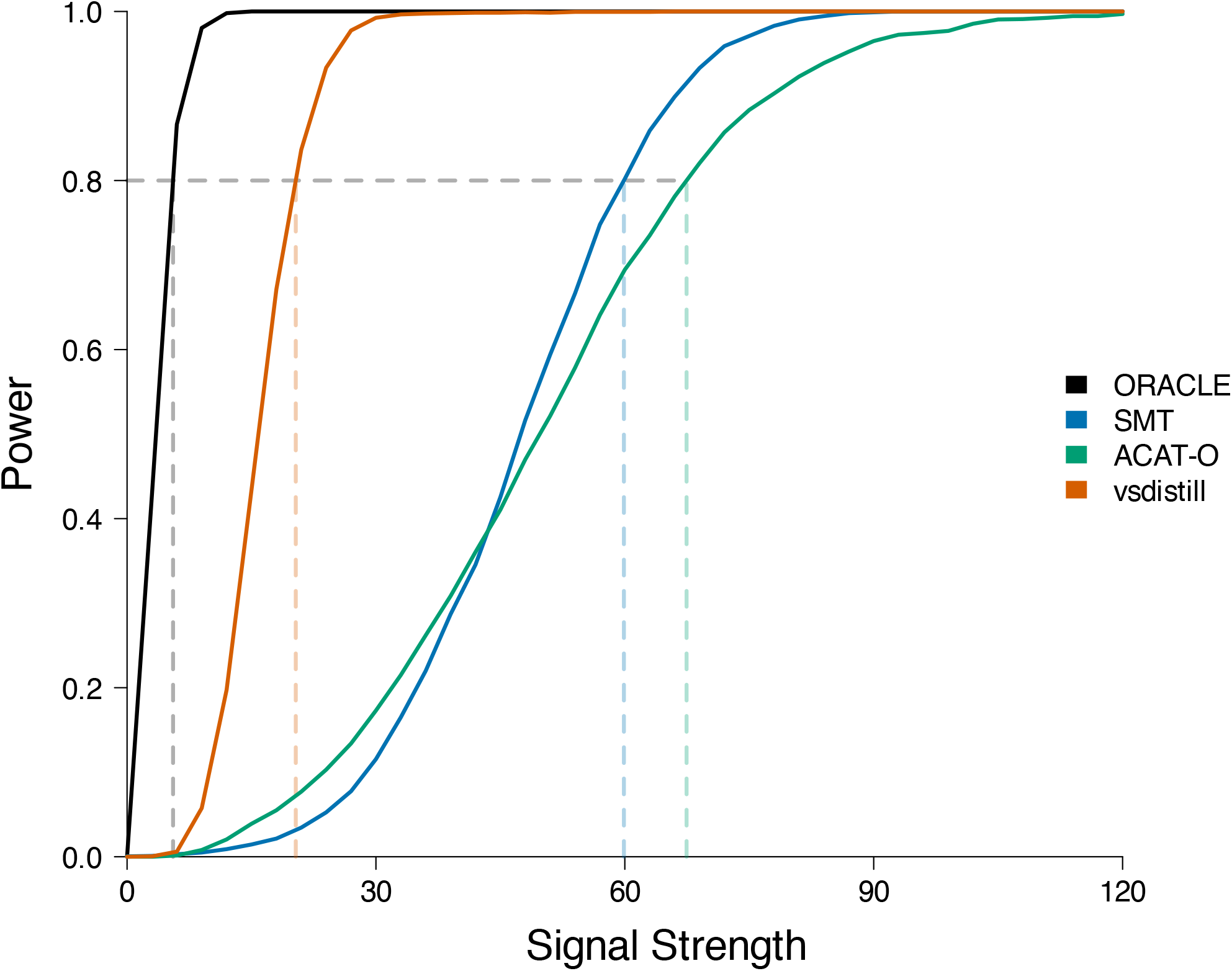
Power comparison between vsdistill, SMT, and ACAT-O in simulations where there are 15 causal variants with equal “expected” effects. The power an oracle ANOVA gene test that knows the causal variants is also included as a benchmark.

**Figure 9.**
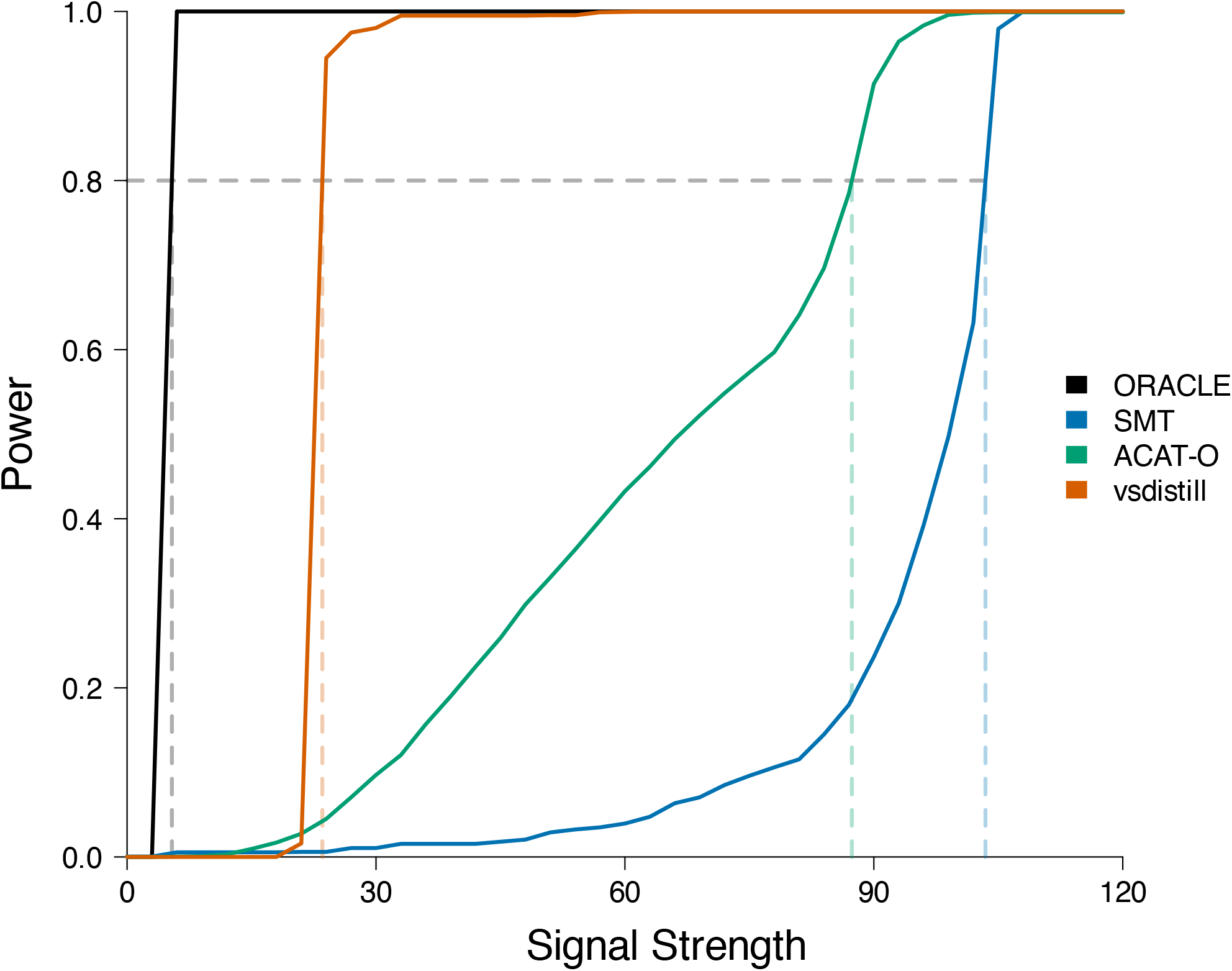
Power comparison between vsdistill, SMT, and ACAT-O in simulations where there are 15 causal variants with equal “obseved” effects. The power an oracle ANOVA gene test that knows the causal variants is also included as a benchmark.

For each gene, we then took 3/2 times the median of the three adjusted p-values as the final p-value (Step 5 in Figure 1). This yielded 451 associations significant at the Bonferroni threshold (*p <* 0.05*/*18, 414). The most significant 10 genes are shown in Table 1. In order to characterize the contribution of rare variation, we repeated our genome-wide screen removing all variants with MAF ≥ 0.01. After calibrating these results with genomic control (*λ* = 1.18, see Figure 3b), we obtained 36 associations significant at the Bonferroni threshold. The most significant 10 genes are shown in Table 2. All of the associations shown in Table 1 and Table 2 have been previously associated with standing height based on the GWAS Catalog (accessed Nov. 14, 2024) [26]. Given the focus of `vsdistill` on leveraging allelic heterogeneity, it is not surprising that *ACAN*, the gene reported to exhibit the highest level of allelic heterogeneity in the genome [35], is found in both tables. Our `vsdistill` implementation reduces memory usage and allocation by storing genotypes, and vectors derived from those genotypes, in a sparse matrix format throughout the procedure. Our implementation also features custom C functions to accelerate our use of this sparse representation and sub-routines to ensure numerical stability. See Section 4.3 and Section 4.4 for details. We ran `vsdistill` across all 18,414 target genes using 250 parallel `mem2_ssd1_v2_x2` instances on the DNANexus Research Analysis Platform (https://ukbiobank.dnanexus.com), each equipped with 2 cores and 8 Gb of RAM. All jobs completed within 1 to 3 hours. Roughly half of the run time was spent on running `vsdistill` in R, the other half was spent reading in genotypes directly from the UK Biobank PLINK exome files. The total genome-wide cost of the compute was approximately $12 USD.

**Table 1.** Leading height genes testing all variants: The 10 genes most significantly associated with height by `vsdistill` based on testing all exonic variants. The − log_10_ p-value is the one reported by `vsdistill` after applying genomic control. Positions are in hg38 coordinates.

| Gene | Entrez ID | Chr | Start Pos. | End Pos. | # Variants | $-\log_{10}(p)$ |
| --- | --- | --- | --- | --- | --- | --- |
| UQCC1 | 55245 | 20 | 35302566 | 35412031 | 201 | 129.96 |
| GDF5-AS1 | 554250 | 20 | 35433029 | 35435450 | 235 | 117.49 |
| GDF5 | 8200 | 20 | 35433347 | 35454746 | 402 | 117.20 |
| LCORL | 254251 | 4 | 17841187 | 18021876 | 2097 | 102.78 |
| ADAMTS17 | 170691 | 15 | 99971437 | 100342005 | 1237 | 98.51 |
| ADAMTS10 | 81794 | 19 | 8580240 | 8610735 | 798 | 79.16 |
| ACAN | 176 | 15 | 88803436 | 88875353 | 1528 | 74.66 |
| CARNS1 | 57571 | 11 | 67415361 | 67425607 | 1081 | 74.06 |
| GNA12 | 2768 | 7 | 2728105 | 2844308 | 603 | 68.28 |
| BLTP3A | 54887 | 6 | 34792083 | 34877514 | 858 | 67.02 |

**Table 2.** Leading height genes testing rare variants: The 10 genes most significantly associated with height by `vsdistill` based on testing exonic variants with MAF *<* 0.01. The − log_10_ p-value is the one reported by `vsdistill` after applying genomic control. Positions are in hg38 coordinates.

| Gene | Entrez ID | Chr | Start Pos. | End Pos. | # Variants | $-\log_{10}(p)$ |
| --- | --- | --- | --- | --- | --- | --- |
| ZFAT | 57623 | 8 | 134477788 | 134832014 | 929 | 34.71 |
| FGFR3 | 2261 | 4 | 1793293 | 1808872 | 838 | 32.01 |
| SCMH1 | 22955 | 1 | 41027202 | 41242306 | 438 | 27.14 |
| ACAN | 176 | 15 | 88803436 | 88875353 | 1516 | 23.68 |
| HAPLN3 | 145864 | 15 | 88877294 | 88895597 | 636 | 21.98 |
| STC2 | 8614 | 5 | 173314723 | 173328447 | 210 | 19.61 |
| GRAMD2A | 196996 | 15 | 72159806 | 72197787 | 197 | 19.23 |
| NPR3 | 4883 | 5 | 32689070 | 32791724 | 483 | 17.73 |
| FBN2 | 2201 | 5 | 128257909 | 128659185 | 1828 | 15.92 |
| CRISPLD2 | 83716 | 16 | 84819985 | 84920768 | 496 | 15.72 |

## 3 Discussion

`vsdistill` provides significant power gains over SMT and ACAT-O in simulation and readily scales to large datasets like the UK Biobank. Future analyses will make use of the ability of `vsdistill` to incorporate prior variant impact predictions and more general gene function predictions, such as those provided by DeepRVAT [9]. We will enable the testing of parallel phenotypes and binary phenotypes with `vsdistill` soon. Existing methods such as STAAR incorporate several important features that are not available in our initial release of `vsdistill`, such as capacity to incorporate close relatives. Extending `vsdistill` to handle these nuanced challenges of genomic analysis will be a focus of future work.

## 4 Methods

### 4.1 The Model

Here we consider testing whether genetic variation within a given variant set, indexed by *𝓁*, modulates a phenotype of interest. Typically, variant sets are defined based on genes or other known functional elements. Let *A* ∈ ℝ^*n*×*q*^ be a matrix of background covariates. Let *K* ∈ ℝ^*n*×*r*^ be a matrix of “high-confidence” predictors that we will always include as part of the alternative model. For example, if putative loss-of-function variant calls are available, *K* may have two columns where the first 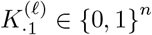 indicates samples that carry at least one putative knock-out variant for gene *𝓁* and the second column 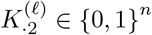 indicates samples that are known to carry a putative knock-out variant on both haplotypes. Alternatively, predictors in *K* may be obtained based on a burden test or using methods such as DeepRVAT [9]. Let 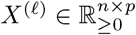 be a matrix of representative hypothesis vectors.

For a set of parameters *α* ∈ ℝ^*q*^, *γ* ∈ ℝ^*r*^, and *β* ∈ ℝ^*p*^, we assume the following model for a quantitative phenotype vector *Y* ∈ ℝ^*n*^.

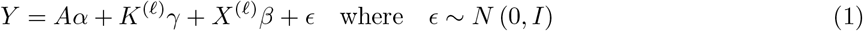

We assume the residuals of *Y* (given *A*) have been rank matched to simulated independent standard Gaussian random variables, so here we take the variance of *ϵ* as known. Under this model we test whether genetic variation at locus *𝓁* affects phenotype *Y* by testing the null hypothesis 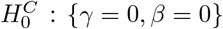 . While our approach is generalizable to binary and survival outcomes, here we only consider quantitative phenotypes.

If *K* is provided, we test *γ* = 0 via the classic likelihood ratio test (a chi-square test) and then treat *K* as part of the background covariates when we continue with testing whether *β* = 0. The p-value returned in this step is later included among the p-values tested by the ROT (Stage 4). In order to properly prioritize the information in *K*, this p-value is given very strong preferential weighting. More explicitly, it is up-weighted to be among the top 5 most significant predictors in *X* (in expectation under 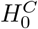).

### 4.2 Hypothesis Generation

Starting with a genotype matrix 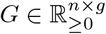 for the *g* variants in our variant set (may be dosages), hypothesis generation maps these genotypes to a matrix of candidate “hypothesis” predictors 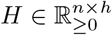. As shown in Figure 1, by default, we simply map each genotype vector to three vectors, each corresponding to a different genetic architecture — recessive, additive, or dominant — as follows.

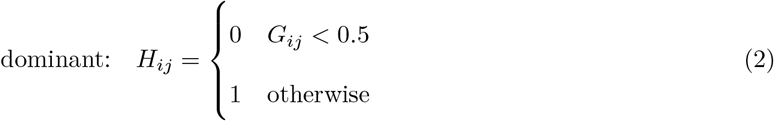

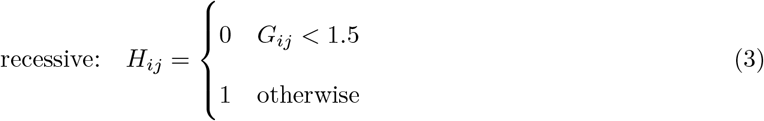

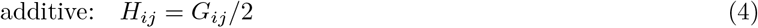

We also assign a prior weight *θ*_*j*_ to each candidate predictor *H*_·*j*_. These weights should be proportional to the probability that the corresponding hypothesis vector has a non-negligible impact on the phenotype of interest. In the absence of any prior information we simply set *θ* = **1**_*h*_. By default, if we only have prior impact prediction weights *w*_*j*_ for each genotype vector *G*_·*j*_, then we set the three entries of *θ* corresponding to *G*_·*j*_ to *w*_*j*_*/*3. Optionally, the genotypes may be interacted with environmental factors, such as smoking status or sex, when constructing *H* in order to generate hypothesis vectors that correspond to pre-determined interaction effects (see Figure 1). We recommend including the base effects for any environmental terms in *A*.

### 4.3 Hypothesis Consolidation

Hypothesis consolidation begins with a matrix *H* ∈ ℝ^*n*×*h*^ where each column corresponds to a candidate predictor and a vector of prior weights 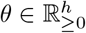 such that *θ*_*j*_ is proportional to the probability that candidate predictor *H*_·*j*_ has a non-negligible effect on the phenotype of interest. If no *θ* is provided, it is set to uniform *θ* = **1**_*h*_ by default. First, we project and normalize each column of *H* to account for our background covariates. More explicitly, given the *QR*-decomposition of (*A, K*) = *QR*, we define 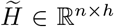 such that 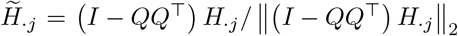. Second, we construct an un-directed graph where each vertex corresponds to a candidate predictor. In this graph, we draw an edge between the vertices corresponding to 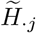 and 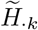 when the observed 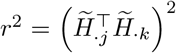 is greater than 0.95. Let *S* ∈ {0, 1}^*h*×*h*^ be the adjacency matrix encoded by this graph. Since this *S* tends to be very sparse, we never explicitly calculate and store 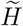 but rather provide a C routine that leverages the sparsity in the original hypothesis matrix *H* to directly obtain *S* in a sparse matrix format.

Finally, we run a greedy clustering algorithm over the graph encoded by *S* to consolidate the candidate predictors into a set of representative predictors. Each round of greedy clustering starts by selecting the vertex *k* with the largest *ψ* = *Sθ* (breaking ties at random). When vertex *k* is selected, *H*_·*k*_ is added as a column to *X* with weight *π* = *ψ*_*k*_. Then, we effectively remove our new cluster from the graph by updating *θ* — assigning 0 to all entries of *θ* such that *S*_·*k*_ == 1 — and the process is reiterated until *ψ* = 0. In the simple case without prior information, *θ* = **1**_*r*_, this approach ensures that the weight *π* assigned to a given column of *X* is proportional to the number of underlying hypotheses (columns of *H*) it represents.

### 4.4 Helical Distillation

Helical Distillation (HD) assigns an independent p-value *p*_*j*_ to each *β*_*j*_ in Equation (1). Here we provide a brief overview of Stable Distillation (SD) sufficient to show how we can build up to HD from a series of SD processes. As above, define 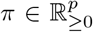 where *π*_*j*_ is taken to be proportional to the prior probability that |*β*_*j*_| is non-negligible. We start by introducing HD in the absence of a pre-defined *π*, in which case the non-informative default is *π* = **1**_*p*_. We will then consider the general case where we start with an arbitrary *π*. A SD process is a stochastic process whereby independent pieces of information relevant to a particular hypothesis are iteratively extracted from a dataset. In this context, we are interested in the null hypothesis *H*_0_ : {*β* = 0}. See Christ et al. [7] for a much more general definition and thorough discussion of SD.

When applied to a high-dimensional regression model like Equation (1), the simple quantile filter based SD procedure presented in Christ et al. [7] involves sequentially iterating over each predictor *X*_·*j*_. Distilling each predictor involves three steps – extraction, filtration, and reconstitution – which can be roughly understood as follows. In the extraction step, we calculate a p-value *p*_*j*_ testing whether *X*_·*j*_ is correlated with the phenotype vector *Y* . In the filtration step, if *p*_*j*_ is smaller than some single pre-specified filtering threshold, then it is stored and used as evidence against *H*_0_; if *p*_*j*_ is larger than that threshold, then a random dummy statistic is stored in its place. In the reconstitution step, a small amount of noise is occasionally injected into *Y* in order to guarantee mutual independence among the *p*_1_, …, *p*_*p*_. This simple SD procedure reports a series of independent p-values, *p*_1_, …, *p*_*p*_, one for each column of *X*.

There is an obvious shortcoming to this simple SD approach: if you start with a matrix *X* where all of the predictors belong to tightly correlated clusters, as expected among nearby genomic variants, then decoupling the predictors and assigning each an independent p-value inflates the multiple testing burden. In other words, testing what is effectively a small number of hypotheses (one for each cluster) as if we were testing many hypotheses (one for each predictor) reduces the power. The hypothesis consolidation step detailed above (see Section 4.3) addresses this issue. The more subtle but fundamental flaw with the simple SD procedure is the requirement to pre-specify a single filtering threshold: in a variant set with allelic heterogeneity, the p-value associated with one causal variant may be orders of magnitude smaller than the p-value associated with another causal variant. In short, there is no “one-size-fits-all” filtering threshold.

HD addresses this problem by iteratively testing each predictor against a series of filtering thresholds while still yielding a single set of p-values *p*_1_, …, *p*_*p*_. HD starts with a SD process that iterates across all predictors with a very stringent p-value threshold. HD then loops through the predictors again, in what we call another “epoch,” this time with a less stringent threshold. Metaphorically, these thresholds form a helical shape as we repeatedly loop over predictors across epochs, hence the name Helical Distillation. The threshold used in the first epoch is selected so that if any of the predictors were to be associated with a p-value below that threshold, we would declare a genome-wide discovery. This makes our HD procedure a function of the number of genome-wide tests the user plans to run in a given experiment. The threshold used in the last epoch is set so that we can be confident that any p-values larger than this threshold will not contribute evidence against *H*_0_. This makes our HD procedure a function of an upper bound *k* on the number of distinct predictors we expect to find among the columns of *X*. In `vsdistill`, we set 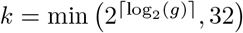 by default. After the last epoch, HD collapses the results across epochs to obtain a single p-value *p*_*j*_ per predictor.

As alluded to above, SD procedures are inherently stochastic: the algorithm must risk introducing a small amount of noise into *Y* at every iteration. Repeatedly testing each predictor for association with *Y* over multiple epochs could potentially amplify this noise, sapping statistical power from our HD procedure. Algorithm 1 employs two novel noise-mitigating features to address this challenge.

Our first innovation: when a significant predictor is “discovered” in the course of distillation, we add that predictor to our background model and continue distilling other predictors conditioned on that updated model. Updating the model for *Y* is justified given the strong Markov property of SD and the fact that updating the model is decidable as a function of p-values that have already been reported by the algorithm. More formally, every time a significant predictor is discovered, we stop the current SD process and reinitialize a new SD process under an augmented model for *Y* . This innovation effectively integrates over the noise that would be introduced to *Y* in a classic SD process after a significant predictor discovery, thereby reducing the stochasticity of the overall algorithm. The fact that this innovation makes our HD process more closely mimic forward stepwise regression supports the idea that it will yield a more powerful approach.

Our second innovation: instead of simply screening for putative p-values below a given upper threshold, in each epoch of HD, we also restrict our search to p-values above a lower threshold based on the upper thresholds used in prior epochs. In other words, when we test a predictor for association with *Y* in a given epoch of SD, we trust that if that predictor was very strongly associated with *Y*, it would have been detected in an earlier epoch. By focusing each epoch on putative p-values within a narrower range, we can significantly reduce the expected amount of noise introduced into *Y* over the course of the HD procedure, thereby conserving power. As further detailed below, in the general case, where we have non-uniform prior weights *π*, we do not use a single upper threshold and lower threshold within each epoch, but rather we offset of the filtering thresholds applied to each *β*_*j*_ based on its corresponding *π*_*j*_ while keeping the average thresholds constant.

For a pre-specified number of *d* epochs, we start with a matrix of upper thresholds *t* ∈ [0, 1]^*d*×*p*^ and a matrix of lower thresholds *s* ∈ [0, 1]^*d*×*p*^. We will revisit how we calculate these thresholds as a function of general prior weights *π* after delineating our HD procedure in the simple case of uniform prior weights *π*. Our HD procedure for the statistical model in Equation (1) is presented in Algorithm 1.

#### Algorithm 1

Helical Distillation for OLS Model with known variance

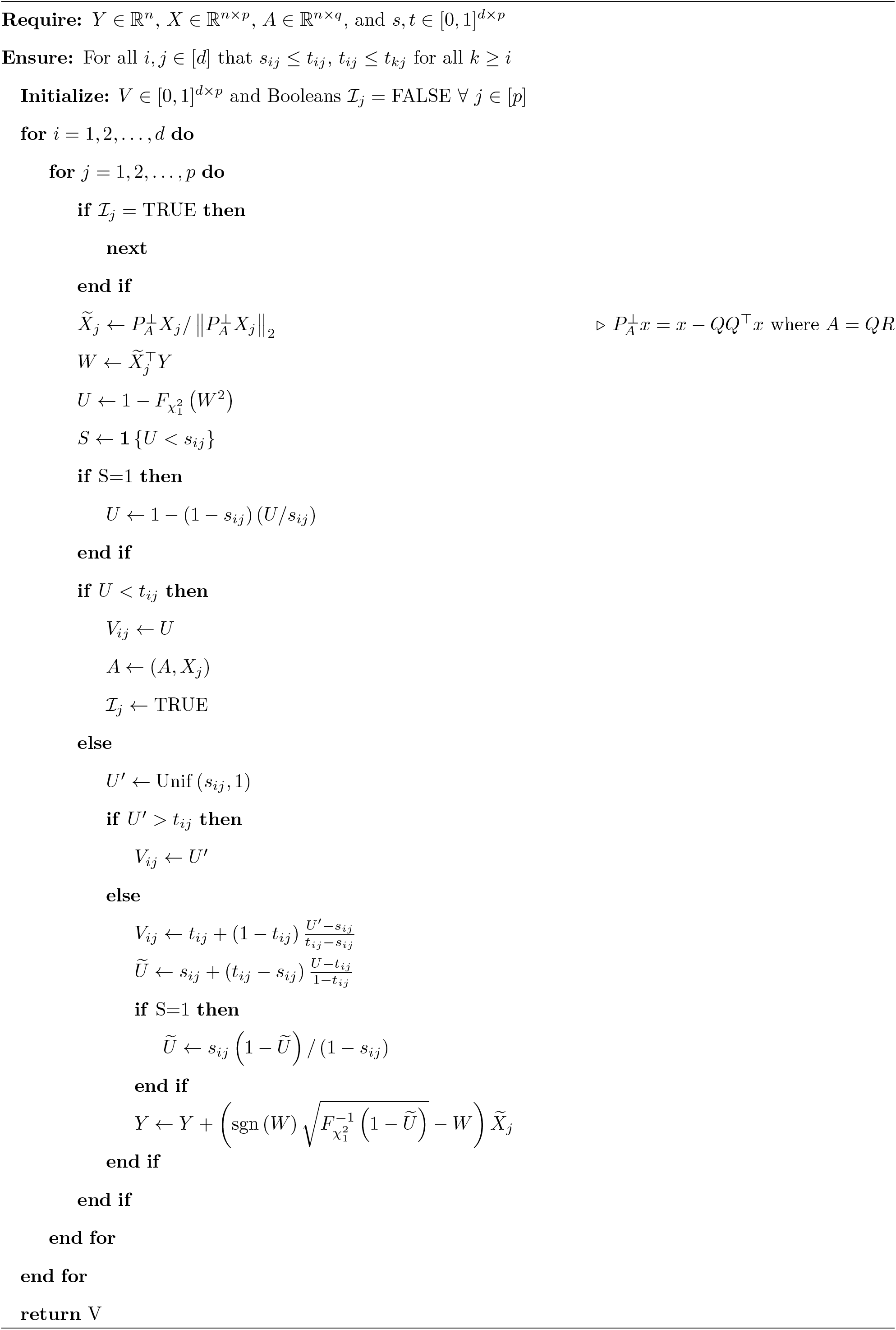

Under the null hypothesis *H*_0_ : *β* = 0, HD guarantees that all entries of *V* are mutually independent where each *V*_*ij*_ ∼ Unif (*s*_*ij*_, 1). While mathematically equivalent, our implementation of Algorithm 1 features several rearrangements and adjustments designed to reduce run time, reduce memory usage, and maintain numerical stability. For example, we compute the inner products required to calculate *W* within each loop with custom C functions that leverage the fact that *X* is typically sparse: *X* has relatively few non-zero entries in genomics applications. Algorithm 1 requires *𝒪* (*npq*) floating point operations (FLOPs); the same as boosting algorithms for ordinary least squares models. While our sparse-*X* implementation only reduces the asmyptotic computational complexity to *𝒪* (*npq/* log(*n*)) FLOPs for real genomic data (based on the expected allele frequency spectrum), in practice the reduction in run time is much more substantial.

Given *V* returned by Algorithm 1, we collapse each column *V*_·*j*_ into a single independent p-value for *β*_*j*_ ≠ 0 using Algorithm 2.

#### Algorithm 2

Collapse V

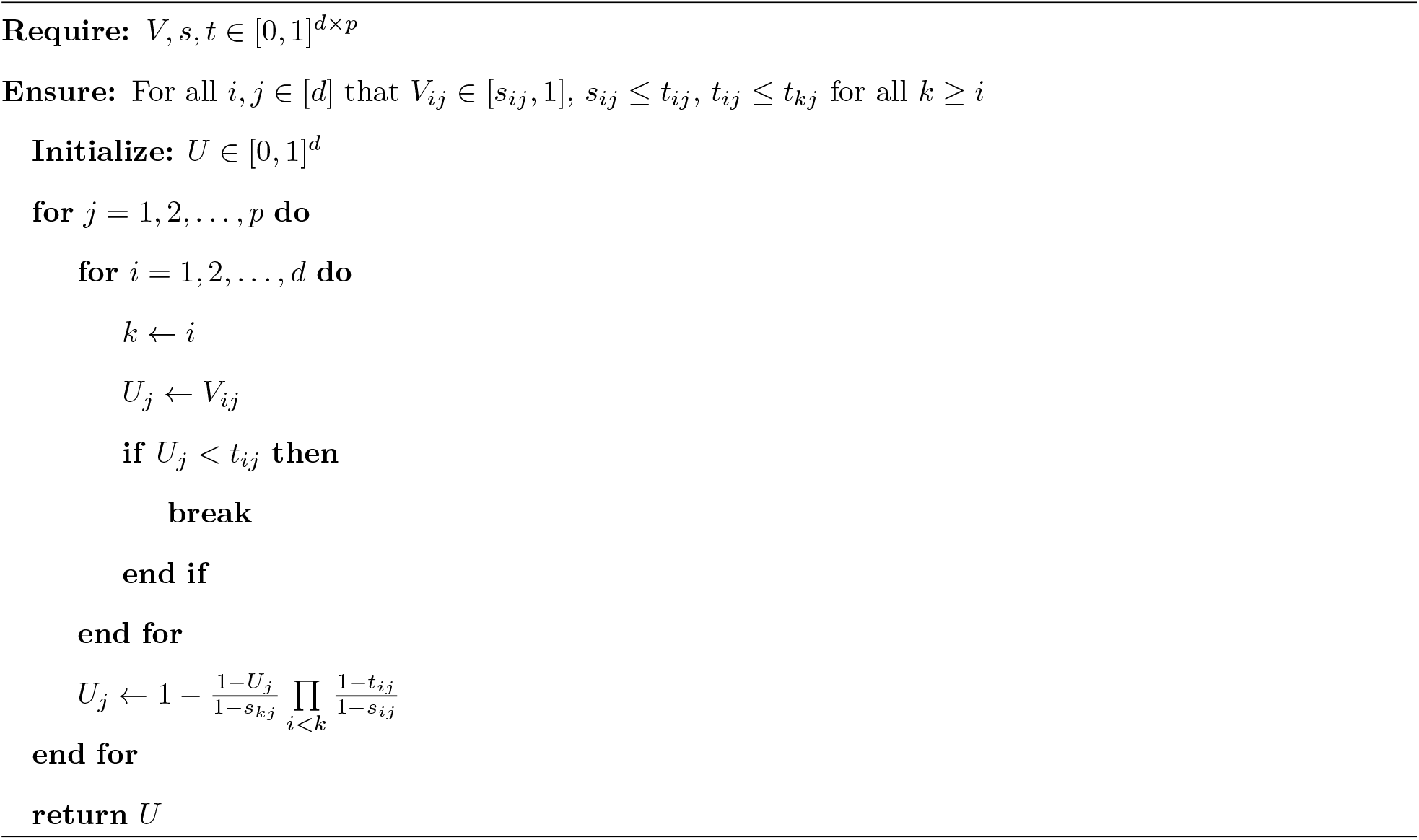

If all entries of *V* are mutually independent where each *V*_*ij*_ ∼ Unif (*s*_*ij*_, 1), then the entries of *U* returned by Algorithm 2 are mutually independent and identically distributed standard uniform random variables. Algorithm 2 is constructed so that any predictor that is discovered to be significant in Algorithm 1 (*V*_*ij*_ *< t*_*ij*_) will returned in the output vector *U* . Thus, this final step of HD captures all of the evidence against *H*_0_ : *β* = 0 in our *d* × *p* dimensional matrix *V* in a single *p*-dimensional vector *U*, as if we had only iterated over the predictors with a single filtering threshold. This collapsing procedure allows HD to avoid paying any significant multiple testing penalty for iterating over the predictors across many epochs.

Now we revisit our choice of threshold matrices, *t, s* ∈ [0, 1]^*d*×*p*^. For all *i* ∈ [*d*] and *j* ∈ [*p*], we require *s*_*ij*_ ≤ *t*_*ij*_ and *t*_*ij*_ ≤ *t*_*kj*_ for all *k* ≥ *i*. Beyond those restrictions, users are free to select *t* and *s* in order to optimize power. Here we present our default specification, which we expect will provide a reasonable power in most applications. We define the threholds we will use in the first epoch of HD, the first row *t*_1·_, based on *T* be our experiment-wide discovery threshold. By default, we set *T* = 0.05*/N*_tests_ given *N*_tests_ conducted genome-wide. We construct a distillation “budget” vector 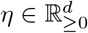 starting with *η*_1_ = *T* . Then, given some multiplier *µ >* 0, we recursively define *η*_*i*+1_ = *µη*_*i*_, terminating when 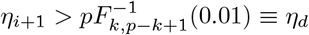 where *F*_*x*,*y*_ is the CDF of the Beta distribution with mean *x/*(*x* + *y*). Note, this process determines the number of epochs *d* we will use in HD. By default, we set *µ* = 4. Given *η* and *π*, we solve for a vector of global thresholds 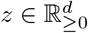 such that each *z*_*i*_ solves the equation

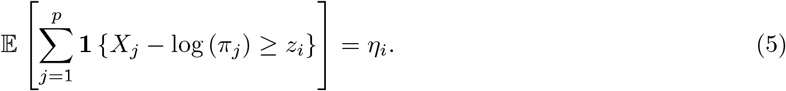

where each 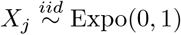. Given *z*, we set *t* _*ij*_ = min (1, exp (−*z*_*i*_ − log (*π*_*j*_))) for all *i* ∈ [*d*] and all *j* ∈ [*p*]. Now turning to *s*, by default we define the first row *s*_1·_ = 0 and each subsequent row as *s*_*i*·_ = 2*t*_*i*−1·_*/µ* for *i* = 2, …, *d*.

The ability to stop and restart the HD procedure based on observed p-values, also enables the dynamic testing of environmental interaction effects (as denoted in Figure 1). If an investigator suspects that, for a particular disease, truly associated variants will tend to have their effects modulated by some environmental variable (eg: smoking status, sex), then upon the “discovery” of a significant predictor, we can reconstitute *Y* and pause the distillation procedure to test the newly “discovered” predictor for interaction effects. As long as these environmental variables and the number of “discovered” predictors we will plan to interact (eg: the first five) are determined *a priori*, then these interaction variables may be treated as if they were initially part of *X* and their p-values used as additional evidence against *H*_0_ by the ROT. This dynamic approach to interaction terms will not be well powered for identifying variants that are only active under one environmental condition (eg: only in smokers); detecting such effects requires adding pre-determined interaction terms to *H*. On the other hand, the dynamic approach avoids much of the additional multiple testing burden required to test pre-determined interaction effects and may improve power for some phenotypes.

## 4.5 Haplotype Data Simulation

In order to assess the calibration and power of `vsdistill`, we simulated 200 genomic datasets, each consisting of a 1 Mb chromosome for 30 thousand human samples (60 thousand haplotypes). Each dataset was simulated using `msprime` [18]. In order to model the diversity of arising genomic datasets, 10 thousand samples in each dataset were drawn from each of three 1000 Genomes populations – Yoruba, Han Chinese, and Central European. We used a demographic model adapted from the `msprime` demography tutorial [12], which itself was based on the population parameters related those three populations presented in [16]. We only made one modification to the demographic model specified in [12]: we use the Discrete Time Wright–Fisher model for the first 100 generations into the past before reverting back to the classic Hudson model as proposed in [24]. The mutation rate was set to 1.2×10^−8^.

Each 1 Mb region of simulated haplotypes was generated using a population-specific human recombination map estimated by `pyrho` [27]. Explicitly, in each simulation, we randomly selected one of our three 1000 Genomes populations (YRI, CHB, or CEU) and 1 Mb segment from genome (excluding heterochromatic regions), then we loaded the recombination map estimated for that population and region by pyrho.

## 4.6 Phenotype Simulation

In the middle of each 1 Mb region, we selected causal variants within a 10 kb causal window. We considered the case of 3, 9, or 15 causal variants. These causal variants were selected uniformly at random from among variants within the 10 kb causal window. Every variant in the 10 kb causal window with derived allele count less than 600 (derived allele frequency less than 0.01) had an equal chance of being selected as a causal variant. If the simulated chromosomes did not include the requisite number of causal variants within the 10 kb causal window, we rejected that simulated dataset and simulated a new set of chromosomes.

Given an active set of causal variants, we simulated *Y* while distributing the observed effects across the causal variants as evenly as possible by manipulating the *QR*-decomposition as done by [7]. Following their approach, let *𝒜* denote the selected set of causal variants and *X*_A_ denote the genotype matrix encoding those causal variants. We set the background covariate matrix *A* ∈ ℝ ^*n*×5^ to *A* = (*e*_1_, *e*_2_, *e*_3_, *a*_1_, *a*_2_) where each vector *e*_1_, *e*_2_, *e*_3_ indicates membership in one of the three simulated sub-populations. We simulated *a*_1_ from independent standard Gaussian random variables and *a*_2_ from independent Rademacher random variables. The background covariate matrix *A* was provided to all testing methods. For all simulations we set the background covariate effects to *α* = (1, −1, 2, −0.5, 0.5)^⊤^. Let *A* = *Q*_*A*_*R*_*A*_ be the QR-decomposition of A. We calculated 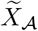 by length-normalizing the columns of 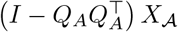. We set 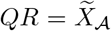 as the *QR*-decomposition of 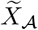.

Given this setup, the sufficient statistic for the oracle ANOVA model that “knows” the causal variants is 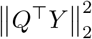 with expected value 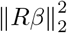. For a desired total association signal strength *s*, we solve 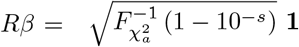 to make the magnitude of each entry of *β* as similar as possible. With *β* now specified for a given *s*, we simulated *ϵ* ∼ *N* (0, *I*_*n*_) and set 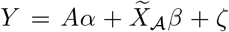. For the equal “expected” signal simulations, we took *ζ* = *ϵ*. The the equal “observed” signal simulations, we took *ζ* = *I* − *QQ*^⊤^ *ϵ*. For the equal “observed” signal simulations, this ensures that the observed 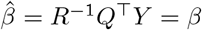 is stable across simulations and that the − log_10_ p-value that one would obtain by testing the resulting *Y* with an oracle model will be approximately *s* in every simulation.

## 4.7 UK Biobank Analysis

### 4.7.1 Variant Selection

We applied quality control to the 470 thousand sample UK Biobank WES genotype data based on the pipeline used by Van Hout et al [29]. Sites with a missing rate greater than 10% or Hardy-Weinberg p-value less than 10^-15^ were removed. We declared HET/HOM genotypes with low coverage (DP<7 for SNV and DP<10 for indels) or HET genotypes with the biased allelic balance (<15% for SNV and <10% for indels) as poor quality using bcftools [11]. We removed sites where the proportion of poor quality genotypes among HET/HOM genotypes was greater than 10%. Additionally, the depth filter recommended by UKBB WES best practices was applied. After variant QC, 20.4 million variants had passed our filters and we used 12.5 million variants in the exome capture regions for our analysis.

### 4.7.2 Sample Selection

We removed samples where reported sex did not match the genetically inferred sex and samples showing sex chromosome aneuploidy or excessive heterozygosity. Samples with a call rate less than 90% were excluded. We then selected samples with European ancestry using UK Biobank data field 22006 (called ‘Caucasian’). After finally removing samples without standing height measurements, we were left with 323,529 samples.

### 4.7.3 Phenotype Testing with vsdistill

A linear model was fit to raw height measurements with the following covariates: sex, age, age^2^, sex x age, sex x age^2^ and 20 principal components provided by the UK Biobank. The residuals of this model were matched by rank to simulated Gaussian random variables to achieve rank-normalization and these values were passed to `vsdistill` . To select our target genes for testing, we excluded genes on the sex chromosomes. After excluding monomorphic variants, genes with fewer than two exonic variants were excluded from our analysis. This left us with a total 18,414 genes for both our initial analysis testing all exonic variants and our follow-up analysis testing only rare (MAF *<* 0.01) variants. All UK Biobank exome analyses were conducted on the Research Analysis Platform (https://ukbiobank.dnanexus.com)

## Acknowledgments

RC thanks the Yale University and Boehringer Ingelheim Biomedical Data Science Fellowship Program for its generous support. This research has been conducted using the UK Biobank Resource under application number 56546.

## Data and Code Availability

`gdistill` will be made publicly available at `ryanchrist.r-universe.dev` shortly. Please to request access to a pre-release version.

## Supplemental Information

